## Supplementary figures and images for "DNA polymerase epsilon is a central coordinator of heterochromatin structure and function in *Arabidopsis*"

### Supplementary figures

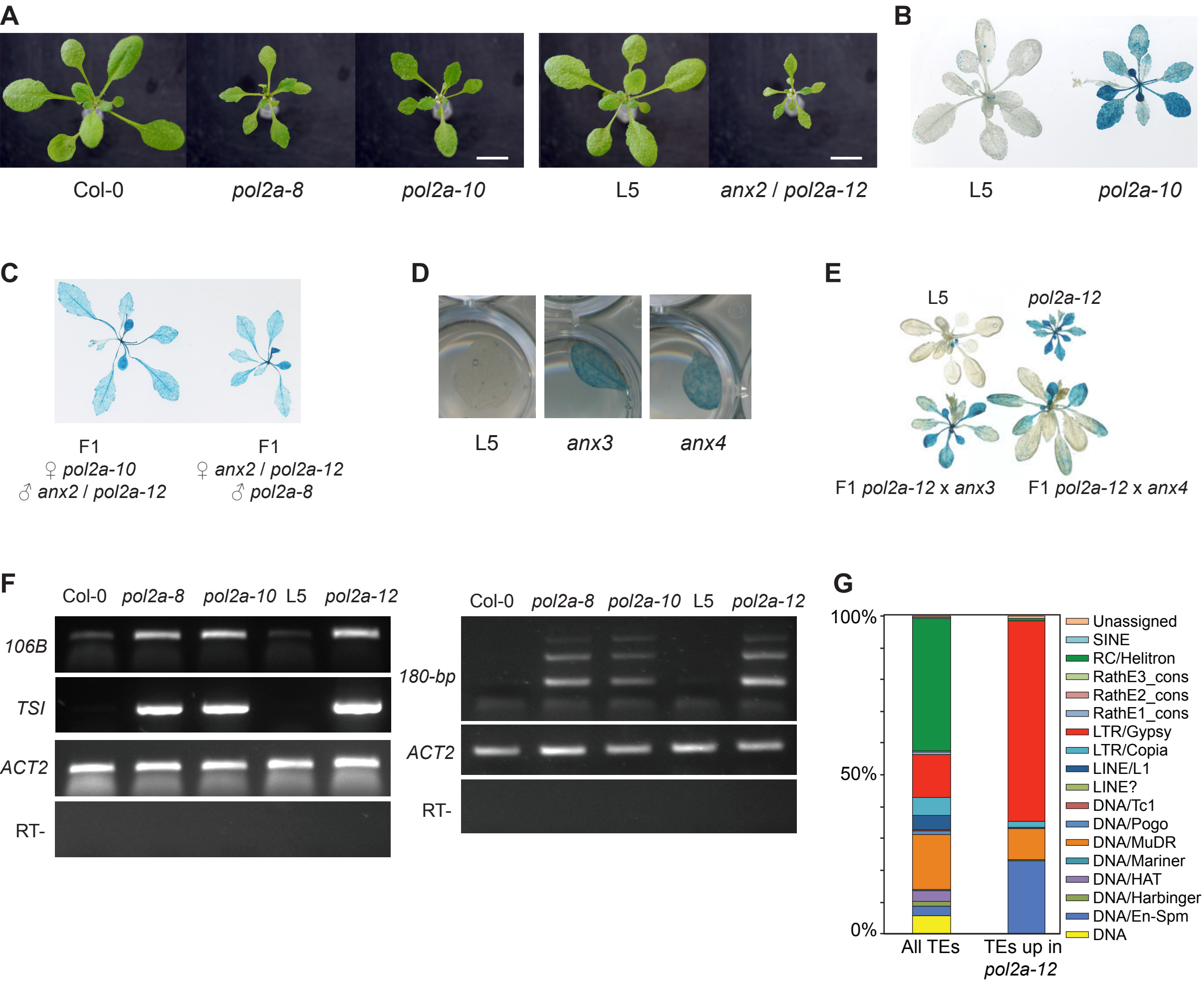

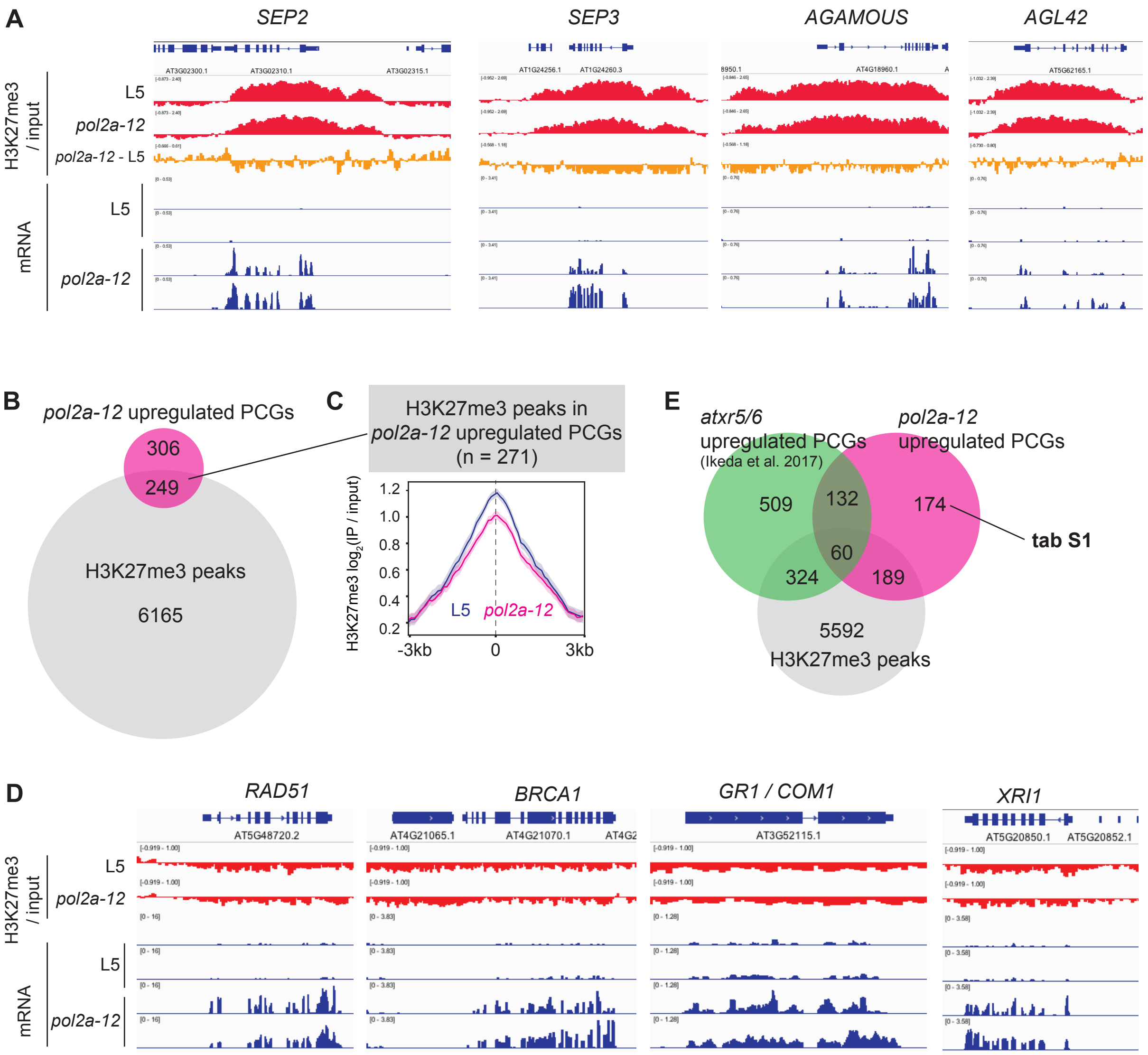

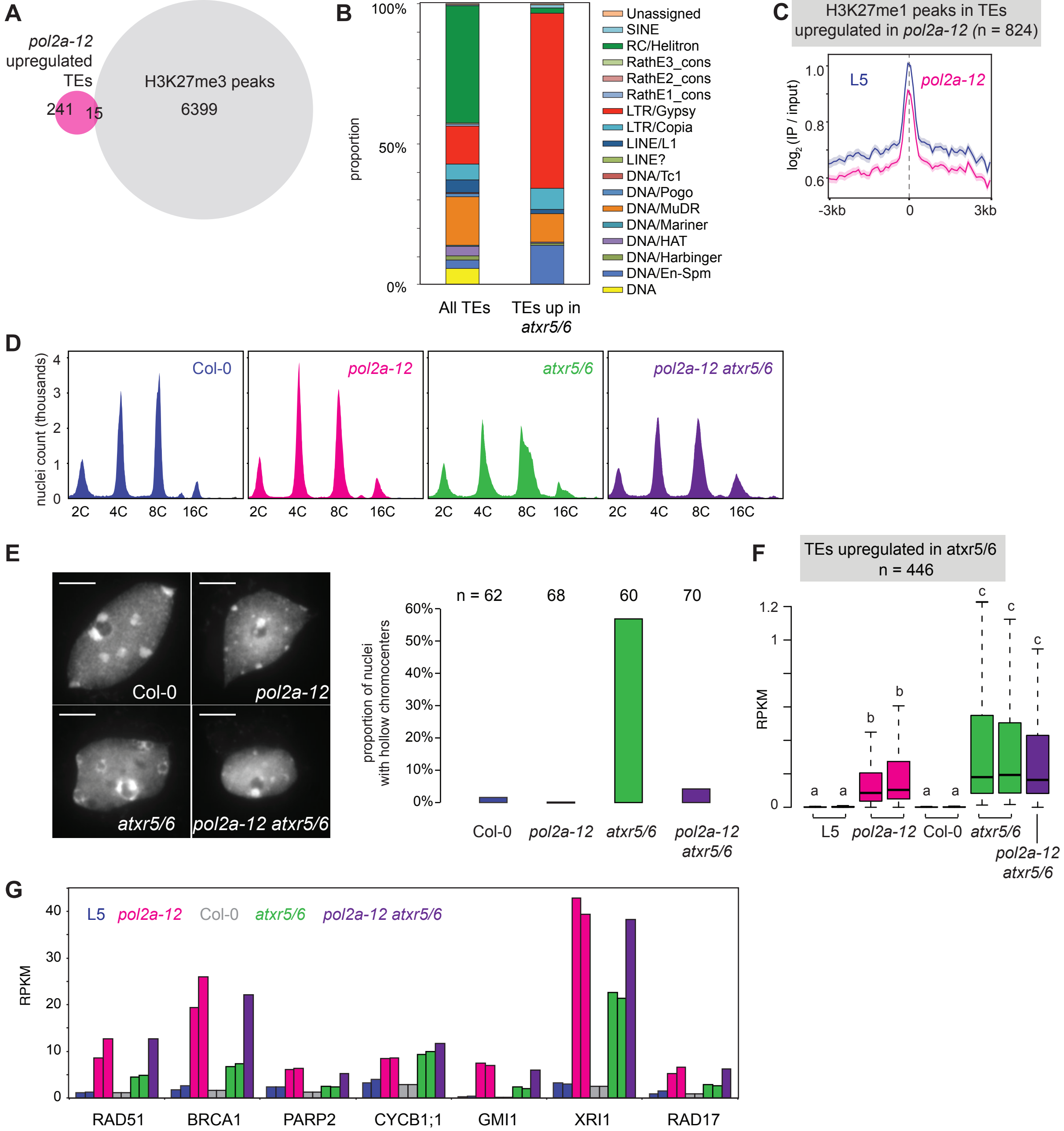

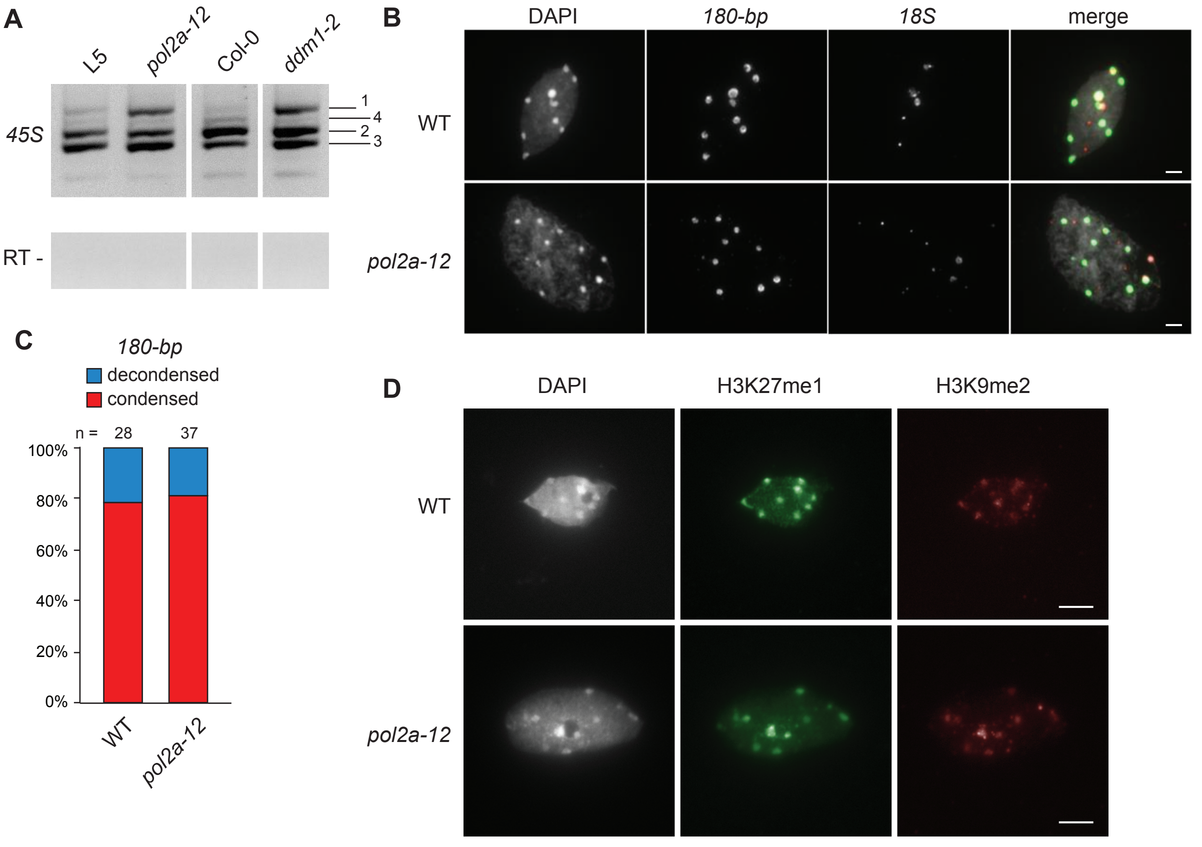

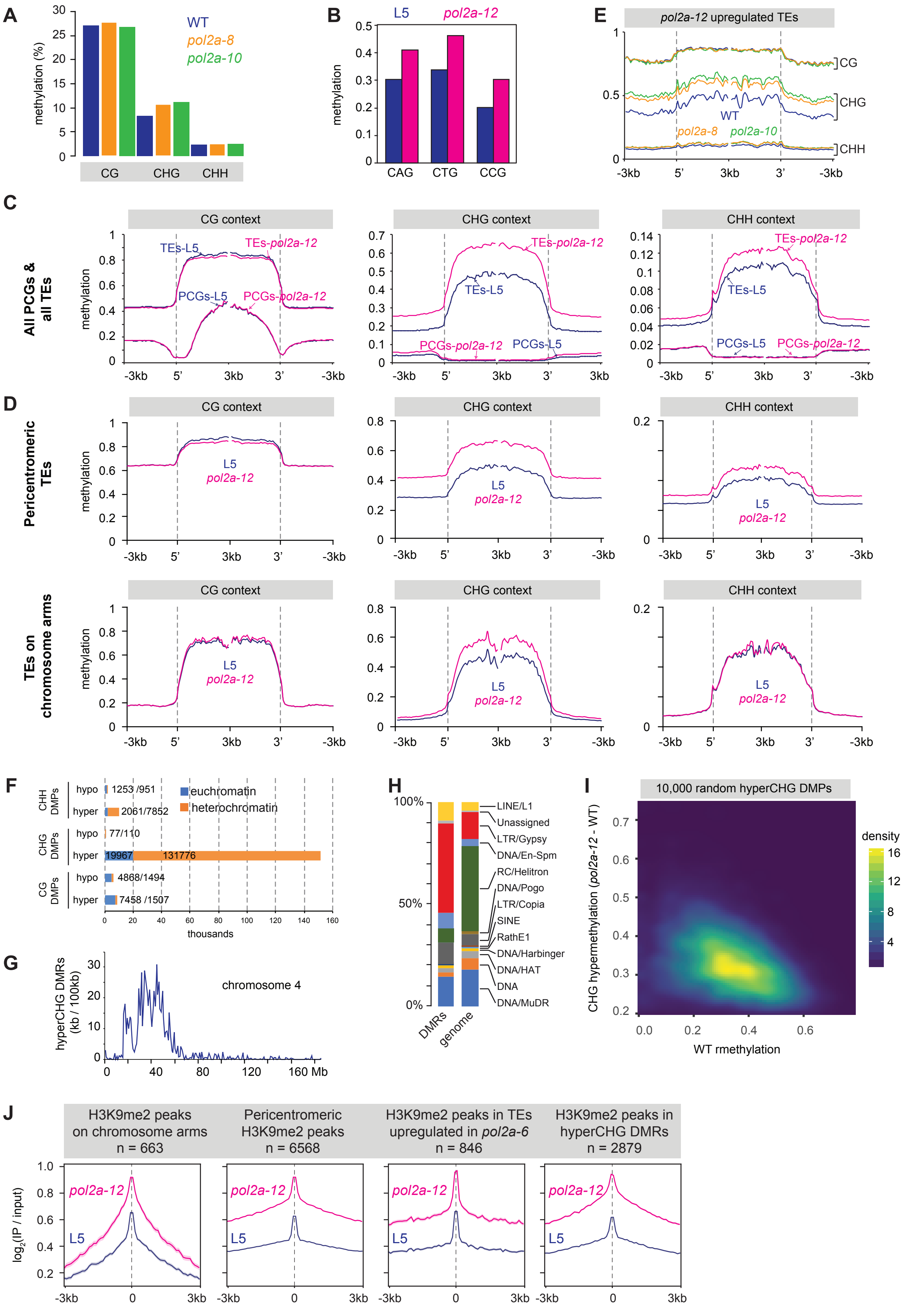

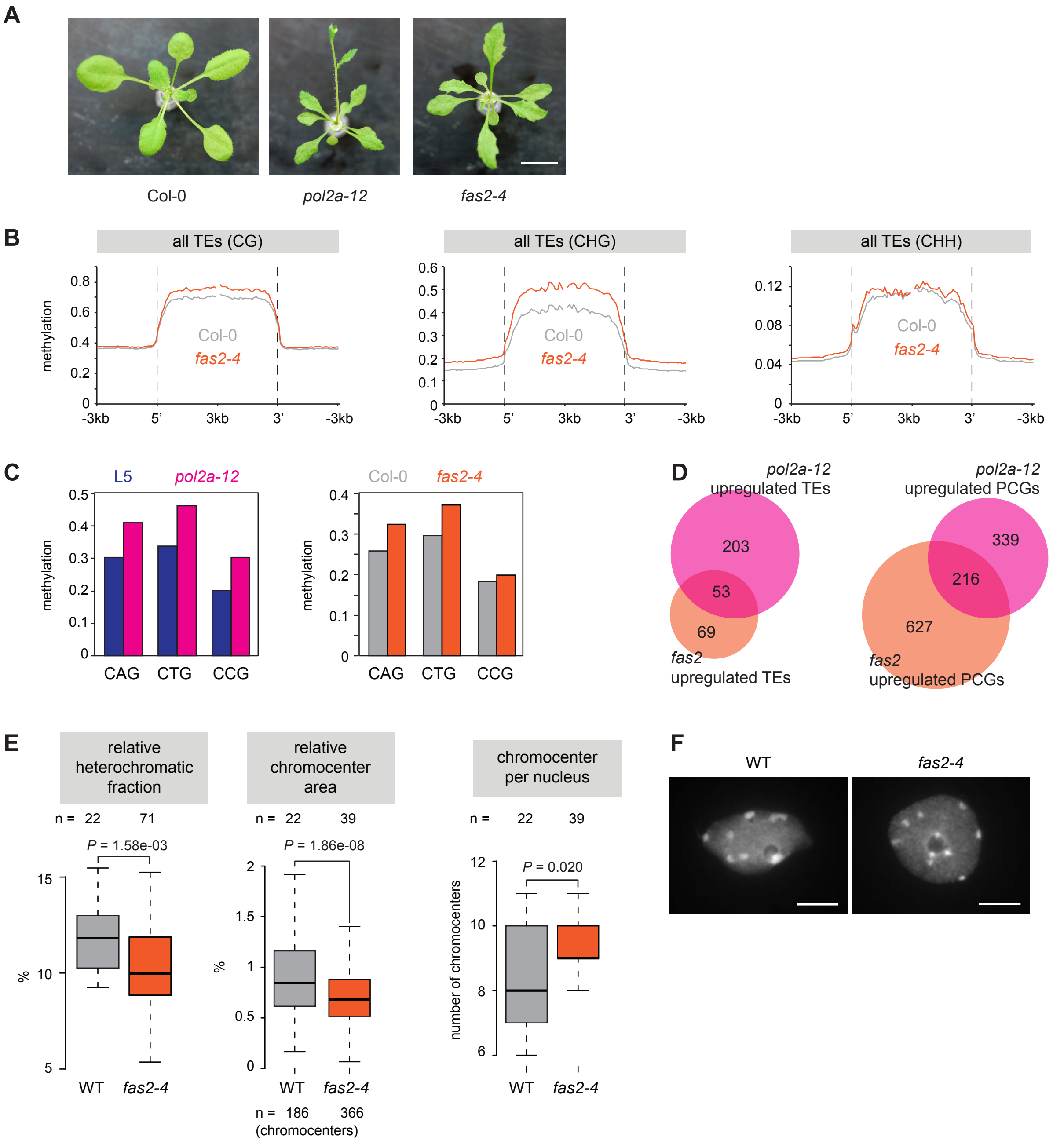

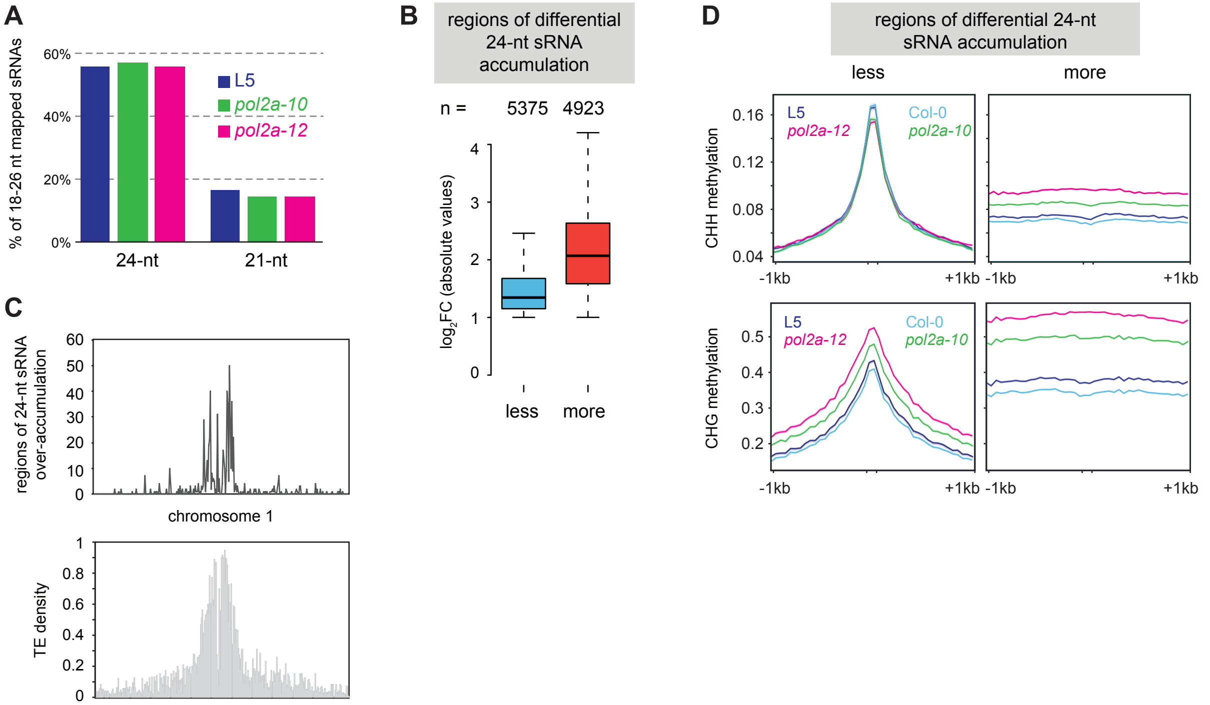

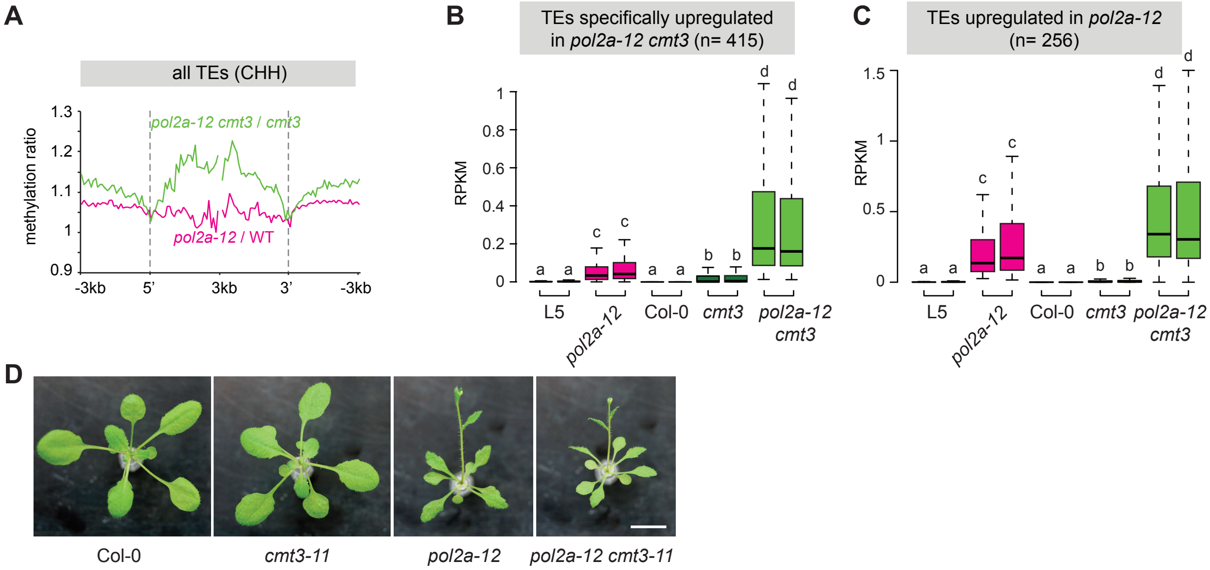

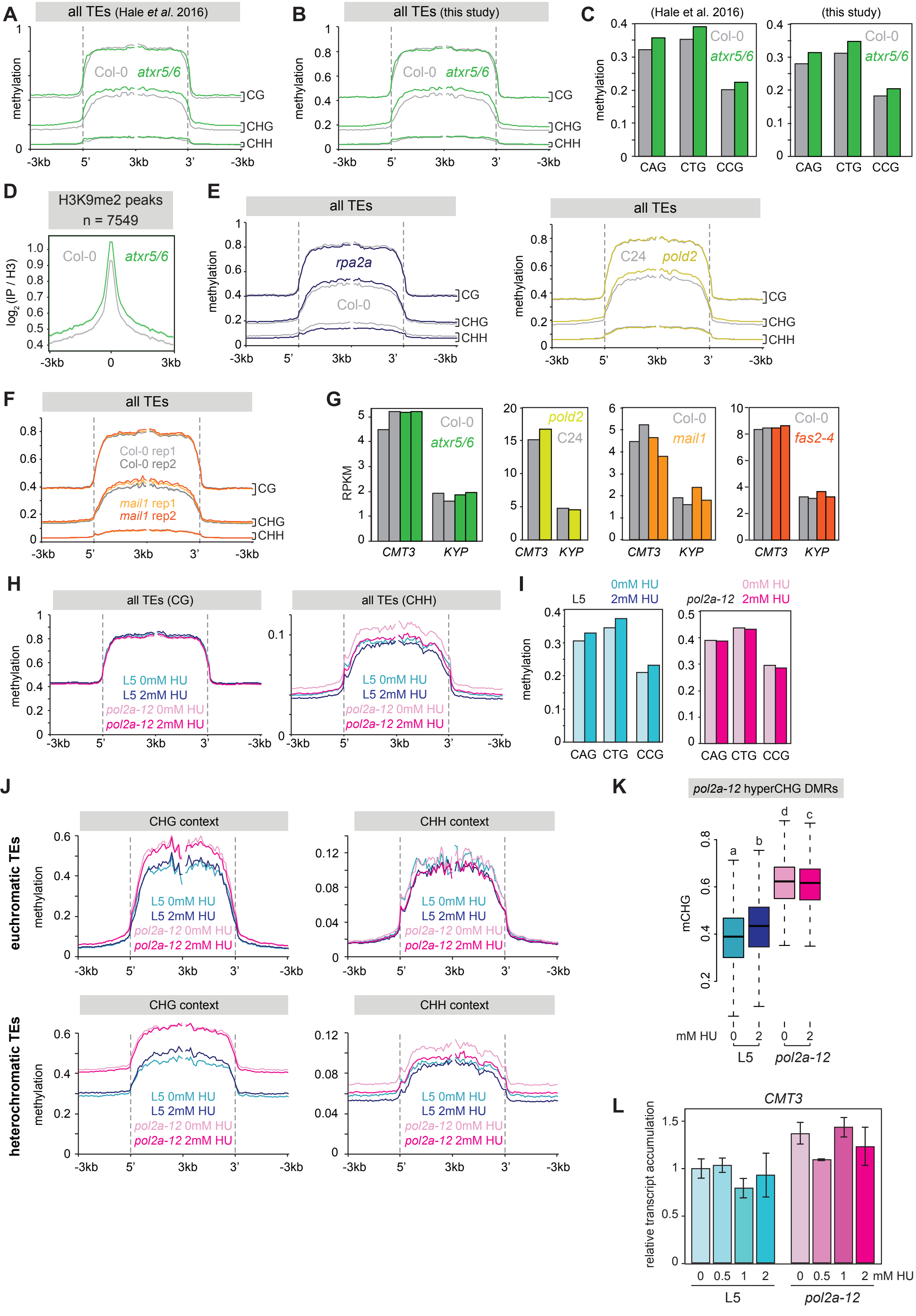
